# A glyphosate-based herbicide cross-selects for antibiotic resistance genes in bacterioplankton communities

**DOI:** 10.1101/2021.12.13.472531

**Authors:** Naíla Barbosa da Costa, Marie-Pier Hébert, Vincent Fugère, Yves Terrat, Gregor F. Fussmann, Andrew Gonzalez, B. Jesse Shapiro

**Affiliations:** Département des sciences biologiques, Université de Montréal, Montreal, Canada; Groupe de Recherche Interuniversitaire en Limnologie et environnement aquatique (GRIL), Montreal, Canada; Department of Biology, McGill University, Montreal, Canada; Département des sciences biologiques, Université du Québec à Montréal, Montreal, Canada; Québec Centre for Biodiversity Science (QCBS), Montreal, Canada; Département des sciences de l’environnement, Université du Québec à Trois-Rivières, Trois-Rivières, Canada; Department of Microbiology and Immunology, McGill University, Montreal, Canada; McGill Genome Centre, McGill University, Montreal, Canada

**Keywords:** Antibiotic resistance genes, indirect selection, herbicide, antibiotic efflux pump, metagenomics

## Abstract

Agrochemicals often contaminate freshwater bodies, affecting microbial communities that underlie aquatic food webs. For example, Roundup, a widely-used glyphosate-based herbicide (GBH), has the potential to indirectly select for antibiotic resistant bacteria. Such cross-selection could occur, for example, if the same genes (*e*.*g*. encoding efflux pumps) confer resistance to both glyphosate and antibiotics. To test for cross-resistance in natural aquatic bacterial communities, we added Roundup to 1,000-L mesocosms filled with water from a pristine lake. Over 57 days, we tracked changes in bacterial communities with shotgun metagenomic sequencing, and annotated metagenome-assembled genomes (MAGs) for the presence of known antibiotic resistance genes (ARGs), plasmids, and resistance mutations in the enzyme targeted by glyphosate (enolpyruvyl-shikimate-3-phosphate synthase; EPSPS). We found that high doses of GBH significantly increased ARG frequency and selected for multidrug efflux pumps in particular. The relative abundance of MAGs after a high dose of GBH was predictable based on the number of ARGs encoded in their genomes (17% of variation explained) and, to a lesser extent, by resistance mutations in EPSPS. Together, these results indicate that GBHs have the potential to cross-select for antibiotic resistance in natural freshwater bacteria.

**IMPORTANCE:** Glyphosate-based herbicides (GBHs) such as Roundup may have the unintended consequence of selecting for antibiotic resistance genes (ARGs), as demonstrated in previous experiments. However, the effects of GBHs on ARGs remains unknown in natural aquatic communities, which are often contaminated with pesticides from agricultural runoff. Moreover, the resistance provided by ARGs compared to canonical mutations in the glyphosate target enzyme, EPSPS, remains unclear. Here we used freshwater mesocosm experiments to show that GBHs strongly select for ARGs, particularly multidrug efflux pumps. These selective effects are evident after just a few days, and at glyphosate concentrations that are high but still within short-term (1-4 day) regulatory limits. The ability of bacteria to survive and thrive after GBH stress was predictable by the number of ARGs in their genomes, and to a lesser extent by mutations in EPSPS. GBHs are therefore likely to select for higher ARG frequencies in natural streams, lakes, and ponds.

## INTRODUCTION

Glyphosate-based herbicides (GBHs) are by far the most extensively used weed-killers worldwide, especially since the introduction of transgenic glyphosate-resistant crops in the 1990s [1,2]. Glyphosate residues can spread widely and accumulate in soil, water, and plant products, raising concerns over human and environmental health [3]. A recent systematic review and risk analysis concluded that glyphosate poses a moderate to high risk to freshwater biodiversity in 20 of the countries investigated [4]. Some of the highest aquatic concentrations of glyphosate were found in countries with the largest production of genetically engineered glyphosate-tolerant crops globally, including the United States, Brazil, and Argentina [2,4].

Although designed to control weed growth, glyphosate may also affect microorganisms that use the herbicide’s molecular target, the enzyme enolpyruvyl-shikimate-3-phosphate synthase (EPSPS), to synthesize aromatic amino acids [5]. The EPSPS is classified into four classes according to mutations in the enzyme active site that confer differential sensitivities to glyphosate [6]. In bacteria, EPSPS classes I and II, which are respectively sensitive and tolerant to glyphosate, are the most frequently found, while classes III and IV are rarer and both confer glyphosate resistance [6]. The EPSPS class II sequence isolated from a strain of *Agrobacterium tumefaciens* is used as the transgene in most commercially available glyphosate-resistant crops [7,8].

Experiments conducted in diverse environments, such as soil and freshwater [9–11] and the bee gut microbiome [12], have shown that bacterial taxa from natural ecosystems vary in their sensitivity to glyphosate. Some of this variation is explained by the distribution of different EPSPS classes. However, while strains with the EPSPS class I are known to be sensitive, they have also been observed to tolerate glyphosate through unknown mechanisms [12], indicating that additional EPSPS-independent glyphosate resistance mechanisms likely exist in nature.

Studies with bacterial cultures have shown an increased resistance to antibiotics after exposure to high concentrations of glyphosate and other herbicides [13–17]. In the presence of glyphosate, the expression of membrane transporters may confer resistance to glyphosate and antibiotics simultaneously [18]. Specifically, multidrug efflux pumps have been experimentally shown to confer resistance to both glyphosate and antibiotics, presumably by exporting a variety of small molecules [13,14, 18]. This is an example of cross-resistance, a mechanism of indirect selection through which one resistance gene or biochemical system confers resistance to other antimicrobial agents [19,20].

Direct selection of antibiotic resistance arises when bacteria are exposed to an antibiotic agent and mutations conferring resistance to this agent are selected [21]. In contrast, indirect selection for antibiotic resistance occurs in the absence of the antibiotic, either via cross- or co-resistance [19,20]. Cross-resistance occurs when the same gene confers resistance to multiple antibiotic agents, while co-resistance occurs when a resistance gene is genetically linked to another gene that is not necessarily an antibiotic resistance gene (ARG), but that is under positive selection.

Most studies of cross-resistance induced by herbicides focused on bacterial isolates in laboratory experiments [13–16,22]. A recent study has shown that herbicide selection increases the prevalence of ARGs in soil bacterial communities, using observational and experimental field data [23]. However, we still lack evidence for aquatic communities, which are of particular interest because herbicides often reach waterbodies through leaching, runoff, and spray drift from agricultural fields [4,24].

Moreover, the extent of direct selection on EPSPS mutations compared to indirect selection on ARGs is unclear. In a previous study, we used 16S ribosomal gene amplicon sequencing to assess how the composition of freshwater bacterioplankton communities respond to a GBH applied alone or in combination with a widely-used neonicotinoid insecticide [11]. As part of the same experiment, we also showed how phytoplankton undergo community rescue in response to lethal GBH doses [25], and how zooplankton community properties were differentially affected by pesticides, even at glyphosate concentrations below North American water quality guidelines [26]. Because GBH was the main driver of changes in the composition of the bacterial community, we expand on our previous work and investigate the effects of the GBH on ARG frequencies in aquatic bacterial communities in this study, using the same outdoor array of experimental ponds (Fig. 1A).

**Fig. 1.**
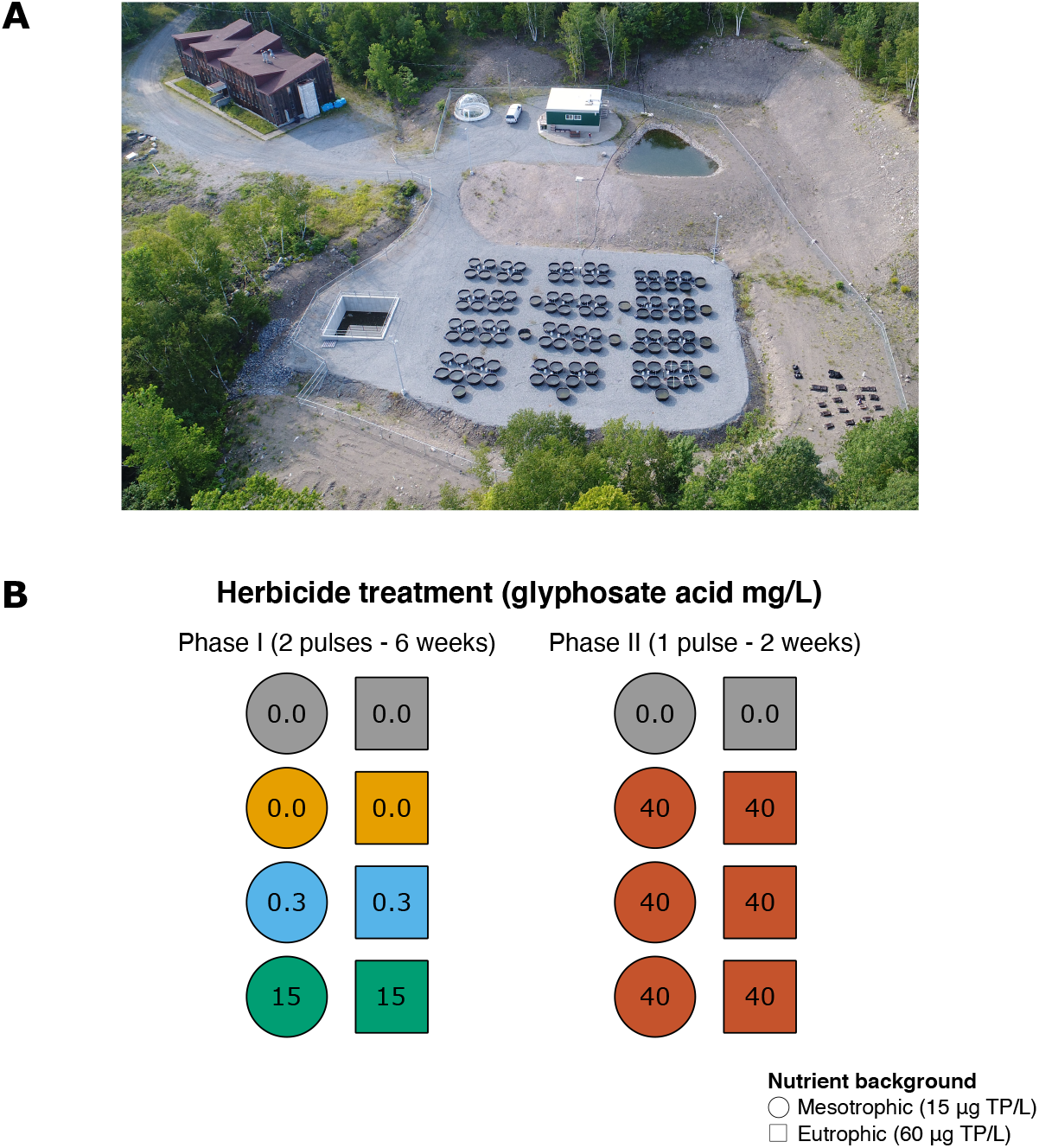
Experimental area and design. (A) Aerial photograph of the Large Experimental Array of Ponds (LEAP) at Gault Nature Reserve, in Mont Saint-Hilaire (Canada). The laboratory facility and inflow reservoir, where water from our source lake was redirected to before filling the mesocosms, can be seen at the top of the photograph. Our source lake, Lake Hertel, is located upstream (not shown in the photograph). (B) Schematic representation of the subset of mesocosms selected for metagenomic sequencing in this study. A total of eight ponds were sampled 11 times over the course of the 8-week experiment, which was divided in two phases: Phase I (6 weeks) and Phase II (2 weeks). Phase I included two pulse applications (doses) of GBH, with three target glyphosate concentrations (0, 0.5, and 15 mg/L). In Phase II, all ponds except for two controls, shown in grey, received a higher dose of glyphosate (40 mg/L). Phase I included four control ponds (grey and yellow) while Phase II only included two controls (grey). Note that yellow ponds only received GBH in Phase II. Nutrients were also added to ponds to reproduce mesotrophic or eutrophic conditions, represented respectively by circles and squares (target phosphorus concentrations are indicated). TP: total phosphorus.

To test the extent to which contamination with GBH cross-selects for ARGs in complex aquatic communities over time, we performed an 8-week experiment in which we exposed freshwater mesocosms to two glyphosate concentrations for six weeks (0.3 and 15 mg/L; Phase I) and to a higher dose for 2 weeks (40 mg/L; Phase II) (Fig. 1B). We sequenced metagenomes from each mesocosm and reconstructed Metagenome-Assembled Genomes (MAGs) of bacteria, which were annotated according to their taxonomy, presence of ARGs, plasmids, and resistance mutations in the EPSPS enzyme. We hypothesize that the frequency of ARGs in bacterial communities increases after exposure to a high concentration of glyphosate, and that efflux pumps are among the main resistance mechanisms promoted by GBH. We also expect that MAGs encoding many ARGs or the resistant classes of the EPSPS gene will be the most likely to survive and proliferate after GBH exposure. Consistent with these expectations, we find that high doses of GBH (15 and 40 mg/L glyphosate) cross-select for ARGs, particularly multidrug efflux pumps. These results show how severe contamination of aquatic systems with GBH could indirectly select for antibiotic resistance.

## RESULTS

### Glyphosate-based herbicide treatment increases antibiotic resistance gene frequency

To test the effects of a GBH on ARGs frequency along the experiment, we tracked variation in the number of metagenomic reads mapped to the Comprehensive Antibiotic Resistance Database (CARD), hereafter referred to as ARG reads, and in the counts of unique ARGs over time, both normalized by the total number of reads in each sample (Fig. 2). In Phase I of the experiment, two pulses of a GBH were applied to reach concentrations of 0.3 mg/L and 15 mg/L glyphosate. Only the latter increased ARG frequencies over time, either when measured as the number of unique ARGs (GAM F=15.65 *p*<0.001, Table 1, Fig. S1), or as the number of ARG reads (GAM F=15.78 *p*<0.001, Table 1, Fig. S1). The concordance of these two metrics suggests that the effect of GBH on ARGs was not due to a few highly responsive resistance genes, but to multiple unique genes. In Phase II, a single dose of 40 mg/L glyphosate was applied to all mesocosms except for the Phase II controls, triggering an increase in ARG frequencies across all treated ponds (Fig. 2). ARG frequencies increased over time, due mainly to the Phase II GBH pulse (Table 1, Fig. S1). Nutrient enrichment produced a weak but significant effect only when considered alone, not in interaction with time (Table 1). Overall, these results support the hypothesis that the GBH treatment has the most dominant and strongest positive effect on ARG frequencies over time.

**Fig. 2.**
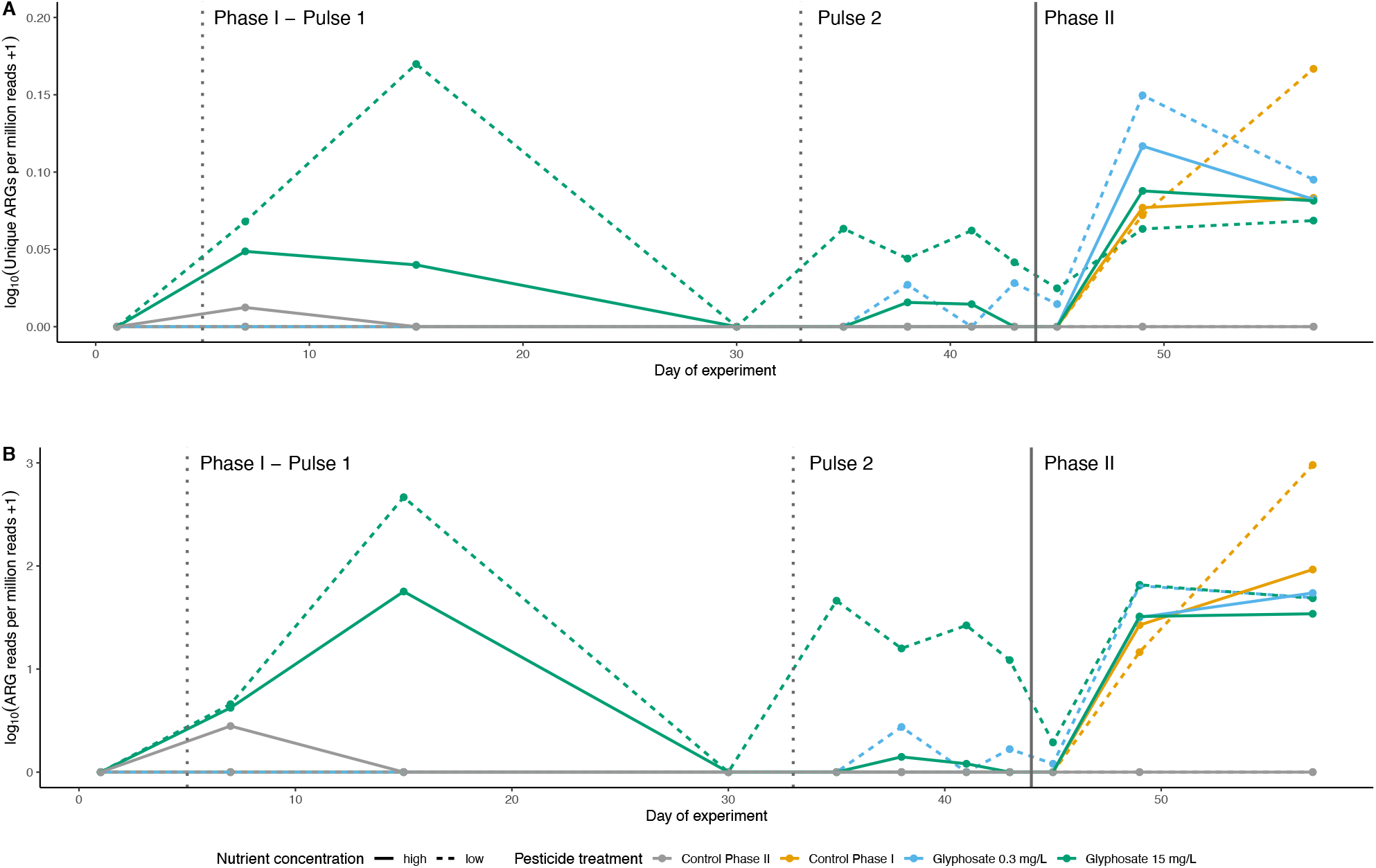
ARG frequencies increase in GBH treatments over time. (A) Number of unique ARGs per million metagenomic reads and (B) number of metagenomic reads mapped to ARGs per million metagenomic reads vary according to treatment and time. Dashed vertical lines indicate the application of Phase I GBH pulses and solid vertical line the Phase II pulse. The colour code refers to the target glyphosate concentrations in Phase I (pulse 1 and pulse 2), while in Phase II all treated ponds received a target of 40 mg/L glyphosate.

**Table 1.**
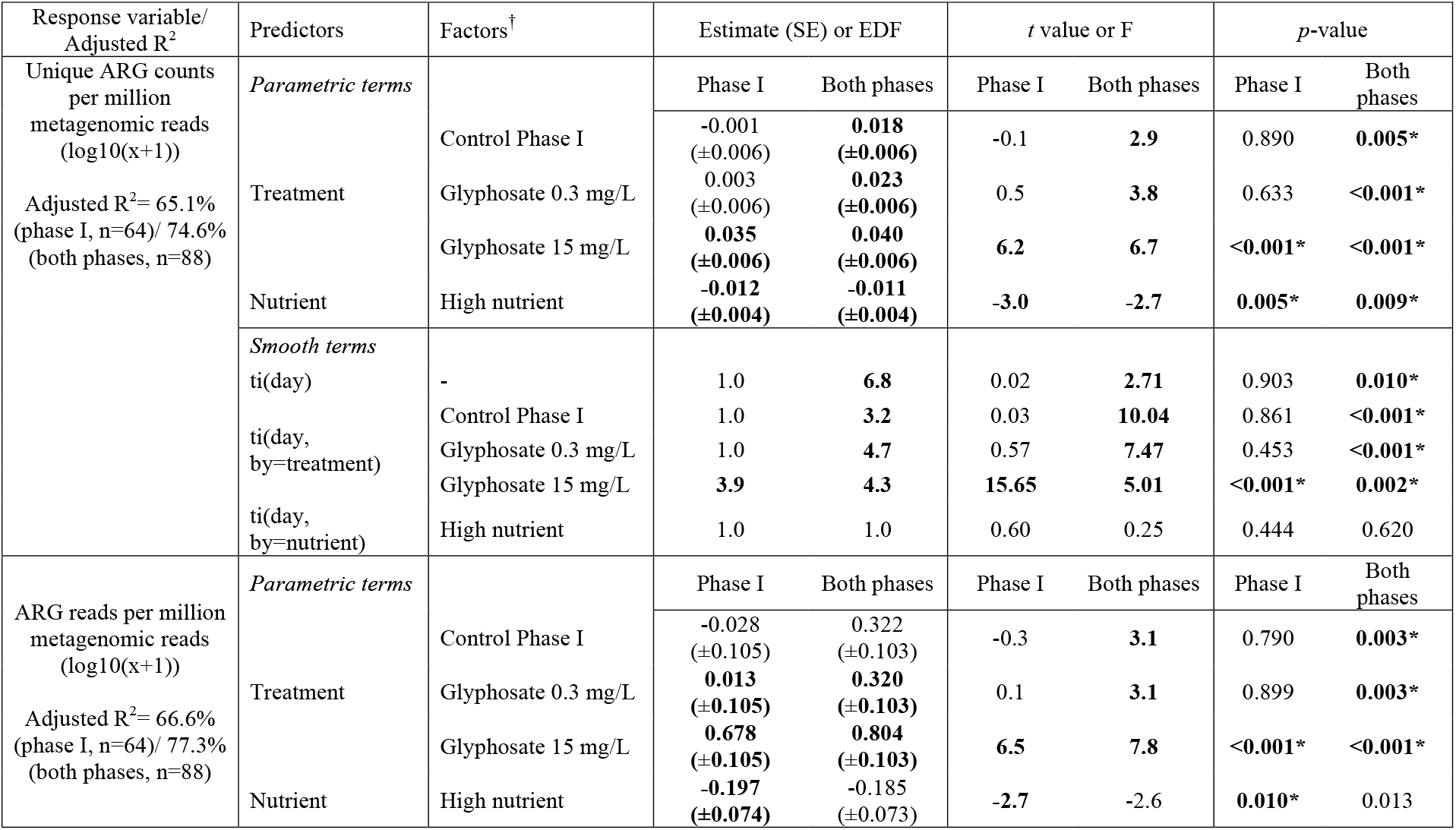
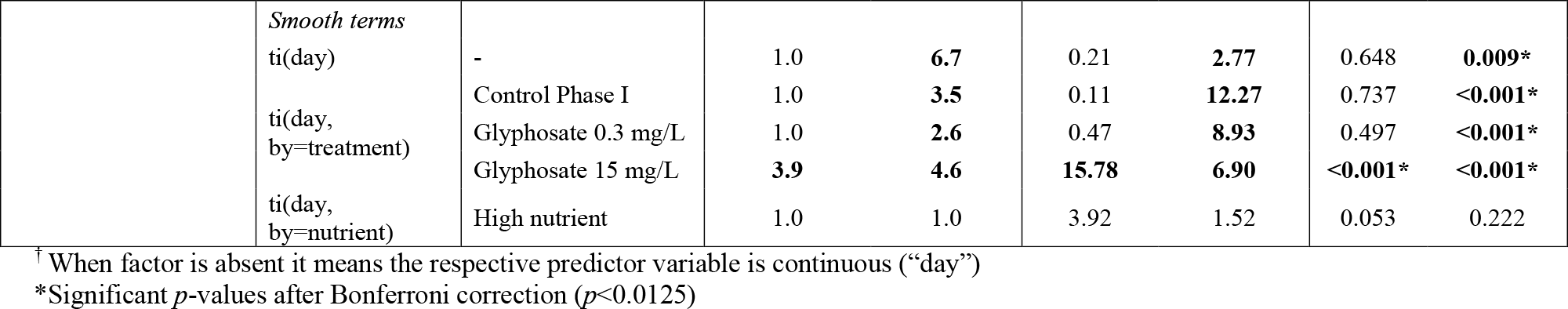
Summary of GAMs showing the effect of GBH on ARG frequencies in phase I only and in both phases. The top rows show unique ARGs as response variable, and the bottom rows show ARG reads. For each predictor of the model, when it is a parametric term we report the respective parameter estimate with standard error (SE) and *t* value. For smooth terms, we report the effective degrees of freedom (EDF) and F statistic. Smooths terms are described as mgcv syntax (‘ti()’ are tensor product interactions). *P*-values are reported for each predictor and reports of significant factors after Bonferroni correction (*p*<0.0125) are highlighted in bold with an asterisk. A Gaussian residual distribution was used.

### GBH selects for specific gene functions, including antibiotic efflux

To assess how GBH affected known gene functions beyond ARGs in the bacterial communities, we built Principal Response Curves (PRCs) based on SEED annotations of genes in the metagenomes. The PRCs revealed a clear effect of GBH on the composition of gene functions (Fig. S2). In Phase I, the first pulse of 15 mg/L glyphosate induced greater deviations from controls than the second pulse. In Phase II, all ponds receiving 40 mg/L glyphosate deviated from the controls. Resistance to antibiotics is among the functions positively affected by GBH treatment, as indicated by the positive scores of the SEED subsystems “Virulence, Disease and Defense”, at level 1 (Fig. S2A), and “Resistance to antibiotics and toxic compounds”, at level 2 (Fig. S2B). Table S1 shows the complete list of PRC scores for all SEED subsystems at levels 1 and 2. Membrane transport (level 1, Fig. S2A), such as the ATP-binding cassette (ABC) transporters (level 2, Fig. S2B), are among the positively selected functions. These genes could plausibly change cell permeability to various molecules, including glyphosate.

To assess the effects of GBH on ARGs at a higher level of resolution, we built another set of PRCs based on ARG profiles predicted from reads mapping to CARD. The resulting PRC plot showed a prominent effect of the first and second pulses of 15 mg/L of glyphosate in Phase I (Fig. 3). In Phase II, the GBH had an effect in all treatments that received a last pulse (40 mg/L glyphosate). This result is consistent with the greater effect of the large Phase II pulse compared to smaller Phase I pulses on total ARG frequencies (Fig. 2 and Fig. S1). The two principal resistance mechanisms of the ARGs annotated by CARD are antibiotic efflux and antibiotic inactivation (shown respectively in blue and red text in Fig. 3). Genes encoding antibiotic efflux functions were more often found with positive PRCs scores (Fisher’s exact test, *p*=0.013), suggesting that they tend to be selected more often than other ARGs in the presence of GBH. This result supports the hypothesis that membrane transporters used for antibiotic efflux could also play a role in exporting glyphosate from bacterial cells.

**Fig. 3.**
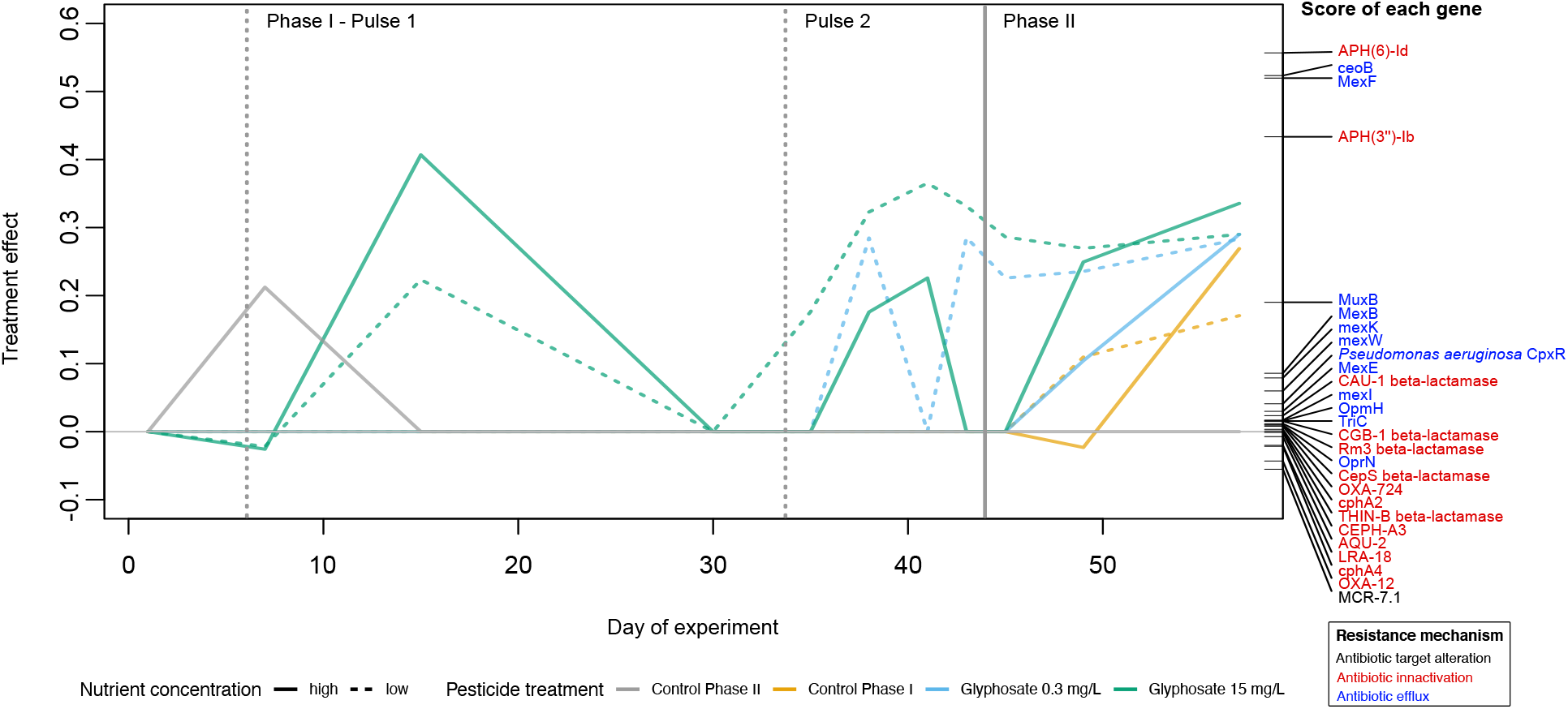
GBH skews composition of ARGs in favour of antibiotic efflux pumps. Principal Response Curves (PRCs) illustrating divergence (relative to controls) in the composition of ARGs in response to GBH exposure. The left y-axis represents the magnitude or ARG compositional response, while the right y-axis represents individual gene scores (i.e., relative contribution to overall compositional changes). Gene names (ARO) are colour-coded based on their mechanism of resistance. Dashed vertical lines indicate the timing of GBH pulses in Phase I, and the solid vertical line represents the pulse in Phase II. The zero line (y=0) represents the low nutrient control pond from both Phase I and II. The PRC explains 30% of the total variance (PERMUTEST, F=22.8, *p*=0.024). Treatments and time interactively explain 74.8% of the variance while 25% is explained by time alone.

### Connecting resistance genes to genomes and plasmids

Thus far, our results have only considered ARGs outside the context of the bacterial genomes or plasmids in which they occur. On average, 71% (± 3; range = 45–94%, Table S2) of ARG reads across samples (those mapping to CARD) also mapped to MAGs, meaning that MAGs captured a large fraction of ARG reads in the metagenomes. We identified putative plasmids in 390 MAGs, with an average of 43 plasmid contigs per MAG (min=1, max=520, SE=3.5, Table S3). However, only 27 plasmid contigs were annotated with ARGs. Out of a total of 188 MAGs with ARGs, only 24 (13%) of them had at least one ARG identified in a potential plasmid. Although some ARGs are certainly encoded on plasmids, ARGs are better associated with genomes than with MAG plasmids in our study.

Of the 426 total MAGs, only 20 recruited 100 or more ARG reads, and the classification of their EPSPS genes varied (Fig. S3, S4). To visualize which ARGs were more abundant in GBH treatments and in which MAGs they were found, we examined the frequency of metagenomic reads mapped to ARGs according to their antibiotic resistance ontology (ARO) classification (top graphs in Fig. S3 and Fig. S4) as well as the proportion of these reads that were mapped to MAGs (bottom graphs in Fig. S3 and Fig. S4). These visualizations confirmed the response of efflux pumps (*e*.*g. mex* genes) to GBH. The relative abundance of *mex* genes is strongly associated with a *Pseudomonas putida* MAG (Fig. S3; bottom right panel) but are sometimes also associated with other MAGs such as *Aeromonas veronii* (Fig. S3), Oxalobacteraceae, and *Azospirillum* (Fig. S4). It is thus likely that GBH selects for efflux pump genes in multiple different genomic backgrounds.

### The number of ARGs encoded in a MAG predicts its frequency after severe GBH exposure

Our results thus far suggest an important role for ARGs, and efflux pumps in particular, in allowing bacterioplankton to survive and grow in the presence of a GBH. We next asked, what is the importance of ARGs relative to genetic variation in the glyphosate target enzyme, EPSPS? Based on known sequence variation in the EPSPS encoding gene, we were able to classify MAGs as putatively glyphosate resistant, sensitive, or unclassified. We also defined a MAG’s antibiotic resistance potential as the number of ARGs identified in their genomes (i.e. number of RGI strict hits). We then tested the extent to which these genomic features were predictive of a MAG’s average relative abundance across ponds at the end of the experiment, after receiving 40 mg/L glyphosate in Phase II. We found that MAGs encoding more unique ARGs tended to have higher relative abundance after receiving the Phase II GBH pulse (Fig. 4A, Table 2). The effect of antibiotic resistance potential was highly significant (multiple linear regression model, *t*=9.53 *p*<0.001, Table 2), and was not observed in control ponds that did not receive the Phase II pulse (Fig. S5A; *t*=2.26 *p*=0.025; not significant after Bonferroni correction, Table S1). The relative abundance of MAGs at the end of the experiment in these control ponds was predicted by their relative abundance in phase I (40% of variance explained; Table S4), consistent with temporal autocorrelation (*e*.*g*. due to random fluctuations in species abundances). In contrast to the strong effect of ARGs on predicting MAG relative abundance post-glyphosate stress (17% of variance explained; Table 2), EPSPS classification explained only 2% of the variation – in both Phase II treatment and control ponds.

**Fig. 4.**
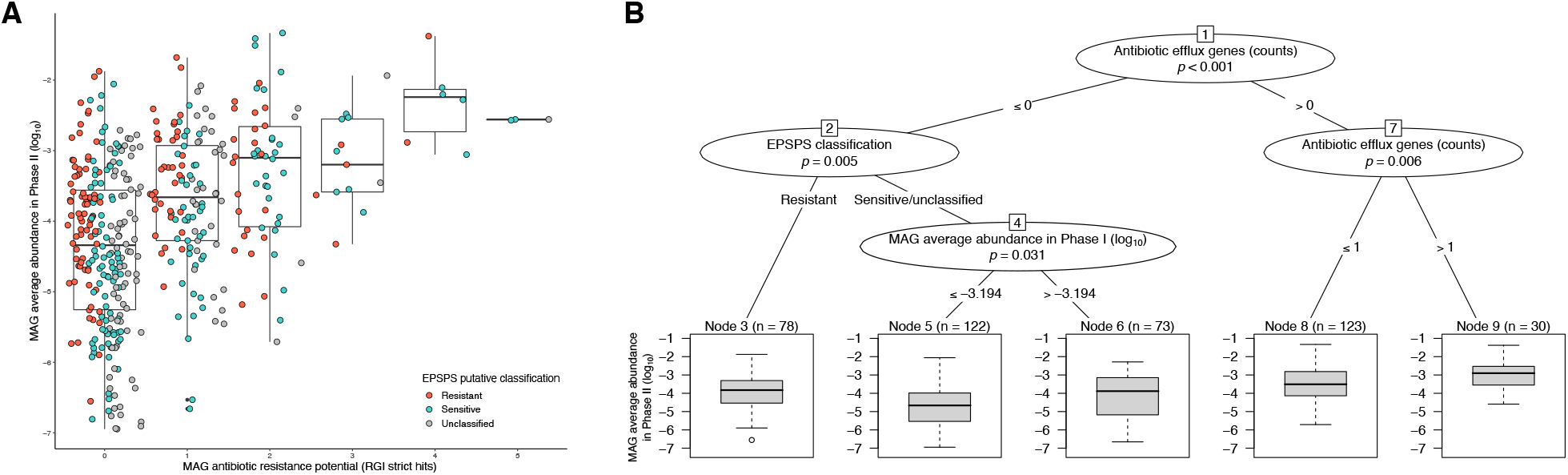
Antibiotic resistance potential predicts MAG relative abundance after severe GBH stress. (A) Boxplots show a positive correlation between MAGs abundance in Phase II and their potential for antibiotic resistance. Each dot represents a MAG that is color-coded based on the predicted resistance of their EPSPS. A slight offset on x-axis (jitter) was introduced to facilitate data visualization. See Table 2 for regression coefficients. (B) Regression tree confirms the significance of the correlation seen in (A), particularly for antibiotic efflux genes. Two other factors were also included, and have small effects on MAG relative abundance in Phase II: the EPSPS classification and the average abundance of MAGs in Phase I.

**Table 2.** Multiple linear regression model and variance partitioning of MAGs abundance in Phase II in treatment mesocosms. *P*-values are reported for each predictor, asterisks indicate significant *p*-values after Bonferroni correction (*p*<0.0125) and reports of significant factors are highlighted in bold with asterisks. Adjusted R-squared equals 21.1% for MAG persistence in treatments (*n*=426, F-statistic: 29.5).

| Response variable | Predictors | Estimate (SE) | *t* value | *p*-value | Explained variance |
| --- | --- | --- | --- | --- | --- |
| MAG mean abundance in Phase II treatment mesocosms | EPSPS classification: |  |  |  | 2% |
| | - Sensitive | 0.002 ($\pm 0.127$) | 0.02 | 0.987 | |
|  | - **Resistant** | **0.413 ($\pm 0.133$)** | **3.11** | **0.002\*** |  |
|  | **MAG antibiotic resistance potential** | **0.496 ($\pm 0.052$)** | **9.53** | **<0.001\*** | 17% |
|  | **MAG mean abundance in Phase I treatment mesocosms ($\log_{10}$)** | **0.178 ($\pm 0.066$)** | **2.69** | **0.007** | 1% |
|  |  |  |  |  | Residuals: 79% |

To further explore these results, we used a regression tree analysis to identify primary drivers of MAG abundance at the end of Phase II. Instead of combining the three major classes of ARGs (antibiotic target alteration, antibiotic inactivation and antibiotic efflux), we used each of them as a separate predictor in the regression tree. The first division splits MAGs with at least one antibiotic efflux gene (Fig. 4B, node 7) which were on average more abundant post-GBH pulse than those without efflux genes (Fig. 4B, node 2). Among MAGs with efflux genes, the more genes they had, the higher their abundance. Among MAGs without antibiotic efflux genes, the EPSPS classification was an important driver of their abundance, followed by the MAG’s average abundance in Phase I. In the absence of a GBH pulse in Phase II, the primary driver of MAG abundance in Phase II controls was their mean relative abundance in Phase I (Fig. S5). Control pond regression trees also included a split between resistant/sensitive and unclassified EPSPS, which is difficult to interpret biologically and likely attributable to noise. This could also explain why 2% of the variation in MAG relative abundance in control ponds was explained by EPSPS class. Together, these results indicate that a bacterial genome’s ARG coding potential is predictive of its ability to persist in the face of GBH stress – more so than the class of EPSPS enzyme it encodes.

## Discussion

Our mesocosm experiment used deep metagenomic sequencing to detect the effect of a GBH Roundup on microbial genes and genomes in semi-natural freshwater bacterial communities. We show that exposure to GBH in high concentrations (15 mg/L and 40 mg/L glyphosate) increases the frequency of ARGs in freshwater bacterioplankton. Moreover, we show that the abundance of MAGs after severe contamination (40 mg/L glyphosate) was predicted based on the number of ARGs they encoded, and these ‘successful’ MAGs tended to have at least one antibiotic efflux gene annotated in their genome. The effect of GBH on ARGs is likely due to cross-resistance, since the multidrug efflux pumps which rise in frequency in response to GBH could potentially transport glyphosate in addition to antibiotics [18]. Alternatively, co-resistance could play a role if GBH selects for bacterial genomes (rather than specific genes) that happen also to encode ARGs. While we cannot exclude a role for co-resistance entirely, the cross-resistance model is more plausible since efflux genes are strongly affected, likely in multiple independent genomic backgrounds. As discussed in detail below, direct selection for EPSPS appears to be weak, implying that ARGs are unlikely to achieve high frequency due to genetic linkage with resistant EPSPS alleles.

An association between glyphosate and increases in ARGs and mobile genetic elements has been previously found in soil microbiomes, as demonstrated in a recent study combining experimental microcosms and environmental data from agricultural field sites in China [23]. Through laboratory assays in three bacterial strains, the authors quantified the conjugation frequency of a multidrug resistance plasmid induced by glyphosate and further investigated changes in cell membrane permeability. They detected a significant increase in conjugation frequency and augmented cell membrane permeability in the presence of glyphosate, suggesting that glyphosate stress increases membrane permeability, thereby promoting plasmid movement. Here, we provide additional support for the hypothesis that cell membrane permeability is altered in the presence of glyphosate, as demonstrated by the selection of membrane transport mechanisms, such as ABC transporters [27] among the annotated gene functions most responsive to the GBH treatments. In contrast, although we did not quantify the frequency of conjugation in our experiment, we did identify some ARGs located on putative plasmids. Of the MAGs encoding ARGs, only 13% contained a plasmid-encoded ARG. It is possible that unassembled plasmids or plasmids not associated with MAGs could harbor ARGs. Including such plasmids would not be expected to change our major conclusion that ARGs are more predictive of MAG frequency post-GBH exposure than EPSPS. In addition to plasmids, other mechanisms also contribute to horizontal gene transfer between bacteria, such as phage-mediated transduction and transformation [28], and future studies could test how these processes may be affected by GBH stress.

Strikingly, antibiotic resistance potential, particularly the presence of antibiotic efflux genes, was more important than the EPSPS classification in explaining variation in MAG abundance in Phase II, after a high GBH pulse. This evidence of cross-resistance in semi-natural communities may help explain why, in previous experiments also performed with complex communities, bacterial strains with the sensitive EPSPS encoding gene were resistant to glyphosate, as it is the case of two strains of *Snodgrassella alvi* in the bee gut microbiome [12]. Although EPSPS alleles were weakly predictive of MAG relative abundance after the phase II GBH pulse, their effects were clearly secondary to the strong effects of ARGs. Computational gene annotations of both ARGs and resistant or sensitive EPSPS have limitations because they are based on sequence similarity, not on phenotypic measurements. Therefore, we cannot entirely exclude a role for EPSPS alleles in conferring GBH resistance in nature, but their effects were small in our experiment. Together, our results strongly suggest that ARGs (and efflux pumps in particular) could be more relevant to glyphosate resistance in nature than mutations in the glyphosate target enzyme.

Our study also aligns with previous single-strain laboratory evidence that antibiotic resistance may enhance bacterial survival in the presence of pesticides. Laboratory assays of bacterial isolates showed accelerated rates of antibiotic resistance selected by exposure to agrochemicals [15,16]. Additionally, it has been shown that the targeted deletion of efflux pump genes can neutralize the increased tolerance to kanamycin and ciprofloxacin in *Escherichia coli* and *Salmonella enterica* serovar Typhimurium in the presence of GBH [13,14]. As we further show in a more natural system, efflux pumps may provide resistance to both glyphosate and certain antibiotics. Whether all efflux pumps are equally capable of transporting various molecules out of the cell remains to be seen, and other resistance mechanisms could also play a role.

It should be noted that we used a commercial Roundup formulation of the herbicide glyphosate, which includes other constituents that may also influence microbial communities and cellular physiology. For example, the surfactant polyethoxylenamine (POEA) has produced negative effects on *Vibrio fischeri* at lower doses than glyphosate acid [29]. However, given that our results are in general agreement with previous soil experiments using pure glyphosate [23], we believe that our findings are at least in part attributable to an effect of glyphosate itself. Furthermore, regardless of whether it is glyphosate or other constituents of GBH that drive cross-selection of ARGs, assessing the risks associated with commercial formulations is more ecologically realistically, as these formulations are used in agriculture fields and lawns [30].

On an applied level, the safety assessment process for pesticides such as glyphosate, currently based on toxicity to model organisms [31,32], should consider the potential effects on bacterioplankton and selection for ARGs. Our results highlight the role of GBH contamination as an indirect selective pressure favouring ARGs in natural communities. Although glyphosate concentrations as high as the ones inducing this effect (i.e. 15 mg/L and 40 mg/L) are rarely found in nature, there are reports of glyphosate levels up to 105 mg/L detected during the rainy season close to agricultural fields; as observed in Argentina [4], for example. Additionally, currently regulated acceptable concentrations of glyphosate in freshwaters in the USA and Canada for short-term exposure (1-4 days) are close to the concentrations used in our experiment (respectively 49.9 mg/L [32] and 27 mg/L [31]). Here we have shown that ARG frequencies can rise dramatically just a few days after GBH treatment, suggesting that even currently acceptable short-term glyphosate exposure could provoke similar selection for ARGs in natural water bodies. The extent to which these ARGs, and the bacteria that encode them, can be mobilized across aquatic ecosystems, and from these ecosystems into animals and humans, remains to be seen.

## METHODS

### Experimental design

An eight-week mesocosm experiment was conducted at the Large Experimental Array of Ponds (LEAP) facility (Fig. 1A) located at McGill University’s Gault Nature Reserve (QC, Canada) from August 17^th^ (day 1) to October 12^th^ (day 57) 2016, as previously described [11,25,26]. Pond mesocosms were filled with 1,000 L of water and planktonic communities from Lake Hertel (45°32’ N, 73°09’ W). Lake water was passed through a coarse sieve to prevent fish introduction, while retaining lake bacterioplankton, zooplankton and phytoplankton, whose responses to experimental treatments have been described in previous studies [11,25,26].

Fig. 1B illustrates the experimental design of a subset of eight treatments selected for the metagenomic sequencing analyses reported here (see [25] for a full description of all treatments at the LEAP facility in 2016). The eight ponds were sampled at 11 timepoints throughout phases I and II of the experiment. In Phase I (days 1-44), all ponds received nutrient inputs biweekly, simulating mesotrophic or eutrophic lake conditions with additions of a concentrated nutrient solution. Four ponds were treated with a GBH to reach target concentrations of 0.3 or 15 mg/L of the active ingredient (glyphosate; acid equivalent), while the other four were kept as control ponds. The GBH was applied in two pulses in Phase I, at days 6 and 33. In Phase II (days 45-57), two control ponds (hereafter referred to as Control Phase I) and the four treatment ponds received one pulse of the GBH at a higher dose (40 mg/L glyphosate) on day 44, while other two other control ponds (hereafter referred to as Control Phase II) received no pulse.

Target doses of the active ingredient were calculated based on the glyphosate acid content in Roundup Grass and Weed Control Super Concentrate (Bayer ©), the formulation used for the experiment. We used a commercial formulation to mimic environmental contamination, and because the costs of using pure glyphosate salt would be prohibitive in a large-scale field experiment. Treatments are referred to by their glyphosate acid concentration to allow comparison with other formulations. Nutrients were added in the form of nitrate (KNO_3_) and phosphate (KH_2_PO_4_ and K_2_PO_4_), with target concentrations of 15 µg P/L and 231 µg N/L in the low-nutrient (mesotrophic) treatment ponds and 60 µg P/L and 924 µg N/L for in the high-nutrient (eutrophic) treatment ponds. The concentrated nutrient solution had an N:P molar ratio of 33 comparable to our source lake. Target doses of glyphosate acid and nutrients were achieved reasonably well, as reported in previous studies [11,25].

### DNA extraction and metagenomic sequencing

The eight experimental ponds were sampled for bacterioplankton DNA at 8 timepoints during Phase I (days 1, 7, 15, 30, 35, 38, 41 and 43) and 3 timepoints during Phase II (days 45, 49 and 57). Water samples were collected with 35 cm long integrated samplers (2.5 cm diameter PVC tubing) at multiple locations in the same pond and stored in 1 L dark Nalgene bottles, at 4 °C until being filtered within 4 hours. We filtered 250 mL of each sample on site, through 0.22 µm pore size Millipore hydrophilic polyethersulfone membranes of 47 mm diameter (Sigma-Aldrich, St. Louis, USA). Filters were stored at -80 °C until DNA extraction.

We extracted DNA from a total of 88 filter samples using the PowerWater DNA Isolation kit (MoBio Technologies Inc.) following the manufacturer’s guidelines. Shotgun metagenomic sequencing was performed using the Illumina HiSeq 4000 technology with 100 bp paired-end reads. Libraries were prepared with 50 ng of DNA using the NEBNext Ultra II DNA Library Prep kit for Illumina (New England Biolabs^®^) as per the manufacturer’s recommendations, and had an average fragment size of 390 bp.

### Metagenomic read trimming, functional annotation and ARGs inference from metagenomic reads

We removed Illumina adapters and quality filtered metagenomic reads using Trimmomatic [33] in the paired-end mode. We used FragGeneScan [34] for gene prediction from trimmed metagenomic reads and annotated predicted genes with SEED subsystems [35]. To identify known ARGs in the metagenomic reads, we used the Resistance Gene Identifier (RGI) ‘bwt’ function that maps FASTQ files of reads passing quality control to CARD [36] using Bowtie2 (version 2.4) as an aligner [37]. Only alignments with mapping quality (MAPQ) higher than 10 and gene coverage of 50% were retained. To calculate the proportion of metagenomic reads mapped to CARD that have been assembled and binned to genomes, we extracted reads that aligned to CARD using Samtools [38] and mapped them to MAGs using Bowtie2 [37]. Table S2 shows the total number of reads by sample after trimming and a summary of the RGI output by sample for hits with minimum gene coverage of 50% and average MAPQ>10.

### Metagenomic *de novo* co-assembly, binning, dereplication and curation of MAGs

We organized the dataset into eight sets of metagenomes, each of them containing samples of the same mesocosm pond (Fig. 1B) from multiple timepoints. We co-assembled reads from each of the 8 timeseries using MEGAHIT v1.1.1 [39], with a minimum contig length of 1 kbp. We used anvi’o v5.1 [40] to profile contigs, to identify genes using Prodigal v2.6.3 [41] and HMMER v3.2.1 [42], to infer the taxonomy of genes with Centrifuge v1.0.4 [43], to map metagenomic reads to contigs using Bowtie2 v2.4.2 [37], and then to estimate depth of read coverage across contigs. Finally, we used anvi’o to cluster contigs according to their sequence composition and coverage across samples with the automatic binning algorithm CONCOCT [44] and we manually refined the bins (*n*=830) using the anvi’o interactive interface, as suggested by developers [40], by removing splits that diverged in the differential coverage and/or tetra-nucleotide frequency of most splits in the same bin.

We dereplicated bins as described in [45]. In summary, we calculated the Pearson correlation coefficient between the relative abundance (i.e. the mean coverage calculated by the function ‘anvi-summarize’ within anvi’o) for each pair of bins in the metagenomic samples, using the ‘cor’ function in R [46], and the average nucleotide identity (ANI) of bins affiliated to the same phylum, using NUCmer [47]. Taxonomy assignment of redundant bins was done using CheckM [48]. Bins with a Person correlation coefficient above 0.9 and ANI of 98% or more were considered redundant. In a total of 830 bins obtained before performing the dereplication, we found 607 non-redundant bins, of which 426 were classified as MAGs, as they had at least 70% completeness and no more than 10% redundancy (see **Table S2**). We then created a non-redundant genomic database of these 426 MAGs to which we mapped metagenomic reads to calculate the relative abundance of each MAGs across the different samples. Here we define a MAG’s relative abundance as the number of metagenomic reads recruited to a MAG divided by the total metagenomic reads in a given sample.

### Identification of ARGs, EPSPS and plasmids in MAGs

We annotated ARGs within MAG contigs with the RGI ‘main’ function, that compares predicted protein sequences from contigs to the CARD protein reference sequence data. Within RGI, we used the BLAST [49] alignment option and the strict algorithm (excluding nudge of loose hits to strict hits) for low quality contigs (<20,000 bp). The RGI low sequence quality option uses Prodigal anonymous mode [41] for the prediction of open reading frames, supporting calls of partial ARGs from short or low quality contigs.

To identify EPSPS sequences from MAG contigs we first used Anvi’o to predict amino acid sequences of the non-redundant MAGs with the flag ‘report-aa-seqs-for-gene-calls’ of the function ‘anvi-summarize’. Gene calls of all the MAGs were concatenated conserving the original split names, and transformed into a fasta file. We then blasted the predicted amino acid sequences against a custom database with sequences of the EPSPS enzyme, using BLASTp [49] and a minimum e-value of 1e-5. After selecting the gene call with the best match (i.e. lowest e-value) to an EPSPS sequence in each of the 426 MAGs, we used the *EPSPSClass* web server [6] to classify the retrieved sequences according to resistance to glyphosate. Sequences were classified as EPSPS class I, class II or class IV if they contained all the amino acid markers from the respective reference, i.e. if the percent identity was equal to 1; and classified as class III when they contained at least one complete motif out of 18 of the resistance-associated sequences, as explained in [6]. MAGs whose EPSPS sequences did not match these criteria of having at least one motif of class III or 100% percent identity with class I, II or IV, or those in which no predicted amino acid sequence matched a known EPSPS sequence were set as unclassified (roughly 27% of MAGs). EPSPS sequences matching class I were considered as putative sensitive and those with at least one motif of class III or matching class II as putative resistant. No sequences were found that matched to class IV.

To identify potential plasmid contigs assembled to MAGs we used the plasmid classifier PlasClass [50]. We counted all contigs classified as plasmid with a minimum of 70% probability, as well as how many of these potential plasmid contigs were annotated with ARGs through RGI. **Table S2** summarizes MAG information, including the predicted EPSPS sequence found in the genome, the EPSPS classification, the number of estimated plasmid contigs and how many of them contained ARG sequences.

### Statistical analyses

All statistical analyses were conducted in R v.4.0.2 [46]. Time series of (log-transformed) ARG counts and ARG reads per million metagenomic reads were modelled using additive models (GAM) using the ‘mgcv’ R package [51]. We used GAMs to account for nonlinear relationships among the response variable and the predictors. Some predictors (nutrient and herbicide treatment levels) were coded as ordered factors; Table 1 lists all factors and predictors of the model. We built the models using the ‘gam’ function and assessed significance of effects with the ‘summary.gam’ function. We validated the models with the ‘gam.check’ function, inspecting the distribution of model residuals, comparing fitted and observed values, and checking if the basis dimension (*k*) of smooth terms were large enough.

We used Principal Response Curves (PRCs) to test for the effect of treatments on the composition of ARGs and gene functional profiles over time. PRCs are a special case of partial redundancy analysis (pRDA) used in temporal experimental studies where treatments and the interaction between treatment and time are used as explanatory variables [52]. Time is the covariable (or conditioning variable) whose effect is partialled out and the response variable is the matrix containing compositional data (taxa or gene family relative abundances). We built PRCs using relative abundances of predicted genes grouped according to the SEED subsystem levels 1 and 2. In a more focused analysis, we built a PRC for the matrix of ARGs found in each sample, i.e. metagenomic reads mapped to each ARG from the CARD reference classified according to their Antibiotic Resistance Ontology (ARO). The matrices were transformed using the Hellinger transformation [53]. The PRC diagram displays the treatment effect on the y-axis, expressed as deviations from the experimental controls at each time point. It also shows species scores on the right y-axis, which here can be interpreted as the contribution of each function or gene to the treatment response curves. We assessed the significance of the first PRC axis by permuting the treatment label of ponds while keeping the temporal order, using the ‘permute’ package [54] followed by a permutation test (999 permutations) using the ‘vegan’ package [55]. For the PRC based on ARG composition, we tested if the distribution of PRC positive and negative scores was different among the resistance mechanisms of the identified ARGs using the ‘fisher.test’ function in the ‘stats’ package in R [46].

To test if MAG abundance in Phase II glyphosate treatments was correlated with their antibiotic resistance potential, we built a multiple linear regression with the ‘lm’ function of the R package ‘stats’ [46]. The response variable was the average relative abundance of a MAG in glyphosate-treated ponds in Phase II. The three predictors were: the MAG’s antibiotic resistance potential (defined as the number of RGI strict hits found in the MAG), the average MAG relative abundance in the same ponds of Phase I, and their EPSPS sequence classification (resistant, sensitive or unclassified). To assess the relative contribution of the different predictors to MAG survival in Phase II, we performed a variance partitioning analysis with the ‘varpart’ function of the R package ‘vegan’ [55]. Finally, to visualize the hierarchy among predictors we constructed a conditional inference regression tree. Response variable and predictors were the same as described above, except that instead of grouping all ARG hits, we transformed them into three variables, according to their function: antibiotic target alteration, antibiotic inactivation, or antibiotic efflux. The regression tree was fitted with the ‘ctree’ function in the R package ‘party’ [46]. As a negative control, we repeated the same analyses for MAGs found in control ponds of Phase II.

As multiple predictors were tested, we performed a Bonferroni correction for the additive and linear models, whereby the *p*-value significance threshold of 0.05 was divided by the number of statistical tests.

Graphs and heatmaps for timeseries data visualization were built using the functions ‘geom_point’ and ‘geom_tile’, respectively, in the R package ‘ggplot2’ [56].

## Data accessibility

Sequence data of the 88 metagenomic samples were submitted to NCBI SRA (BioProject PRJNA767443, accession numbers SRR16126824-SRR16126911) and the genomes of 426 predicted MAGs have been deposited and associated to the same BioProject (BioSample accession numbers in Table S3). The data will be publicly available once the manuscript is accepted for publication, and it can be now accessed through the following reviewer link: https://dataview.ncbi.nlm.nih.gov/object/PRJNA767443?reviewer=vk9o7uf95h5cm1d8m1mqnrc5fk.

## Acknowledgements

We are grateful to D. Maneli, C. Normandin, A. Arkilanian and T. Jagadeesh for their assistance in the field, to J. Marleau and O.M. Pérez-Carrascal for their assistance in the laboratory and to O.M. Pérez-Carrascal for the advice on bioinformatic analyses.

## Funding information

This study was supported by a Canada Research Chair and NSERC Discovery Grant to B.J.S. N.B.C. was funded by FRQNT and NSERC-CREATE/GRIL fellowships. M-P.H. was funded by NSERC and NSERC-CREATE/GRIL. V.F. was supported by an NSERC postdoctoral fellowship. LEAP was built and operated with funds from a CFI Leaders Opportunity Fund, NSERC Discovery Grant and the Liber Ero Chair to A.G.

## Supplementary Material

### Tables

**Table S1** PRC scores from functional annotations shown in Fig. S2

**Table S2** Metagenomic sample information, summary of RGI output for hits above mapping threshold (MAPQ>10 and minimum of 50 gene percent coverage) and proportion of sample reads mapped to CARD (ARG reads) that mapped back to MAGs.

**Table S3** MAG information, predicted EPSPS amino acid sequence, summary of ARGs and plasmids. For each predicted EPSPS sequence, the putative classification regarding glyphosate resistance is shown. The number of potential plasmid contigs and how many of these had ARGs annotated is also shown. Number of ARGs annotated to MAG contigs (total RGI strict hits) are provided in the last column.

**Table S4** Multiple linear regression model and variance partitioning of MAGs abundance in Phase II in control mesocosms. P-values are reported for each predictor, asterisks indicate significant p-values after Bonferroni correction (p<0.0125) and reports of significant factors are highlighted in bold. Adjusted R-squared equals 43.2 % for MAG abundance in controls as response variable (n=425, F-statistic: 78.7).

### Supplementary figures

**Fig. S1 Glyphosate increases ARG frequencies in experimental ponds**. GAMs illustrating the time-dependent effect of GBH and nutrient treatments on unique ARGs in Phase I (A), in both Phase I and II (B), on ARG reads in Phase I (C), in both Phase I and Phase II (D). Dashed vertical lines indicate the application of Phase I GBH pulses and solid vertical line the Phase II pulse. Glyphosate acid concentration of pulses applied in Phase I (dose 1 and dose 2) are indicated in the legend, while in Phase II, all treatments received 40 mg/L, except the Control Phase II. Shades indicate a confidence interval of 95%.

**Fig. S2 Principal Response Curves of the experimental treatment effect on the composition of gene functional profiles predicted from metagenomic reads grouped according to (A) SEED subsystem level 1 and (B) level 2**. Treatment effect is shown in the left y-axis while scores of genes (proportional to their contribution to the treatment effect) are shown in the right y-axis. Dashed vertical lines indicate the application of Phase I glyphosate pulses and solid vertical line the Phase II glyphosate pulse. Glyphosate concentration of pulses applied in Phase I (dose 1 and dose 2) are indicated by the legend, while in Phase II all treatments received 40 mg/L of glyphosate, except the Phase II controls. Treatment effect zero is equivalent to the low nutrient control Phase II pond. Function of resistance to antibiotics is highlighted in red according to how it is named in (A) SEED subsystem level 1 (50.9% of total variance explained, PERMUTEST F=43.1 *p*=0.023) and (B) SEED subsystem level 2 (33.1% of total variance explained, PERMUTEST F=25.8 *p*=0.027), where only scores with absolute values larger than 0.05 are reported (all scores are shown in Table S1).

**Fig. S3 Metagenomic reads mapped to ARGs classified according to their ARO (top graph) and ARG reads mapped to MAGs (bottom graph) in low nutrient ponds**. MAG identities are followed by their finest taxonomic assignment (o=order, f=family, g=genus, s=species). Only alignments with MAPQ>10 were tallied. Dashed vertical lines represent Phase I GBH and solid vertical lines are Phase II pulses (all at 40 mg/L glyphosate).

**Fig. S4 Metagenomic reads mapped to ARGs classified according to their ARO (top graph) and ARG reads mapped to MAGs (bottom graph) in high nutrient ponds**. MAG identities are followed by their finest taxonomic assignment (o=order, f=family, g=genus, s=species). Only alignments with MAPQ>10 were tallied. Dashed vertical lines represent Phase I GBH pulses and solid vertical lines are Phase II pulses (all at 40 mg/L glyphosate).

**Fig. S5 MAG mean relative abundance in controls of Phase II as a function of antibiotic resistance potential (or the amount of ARGs annotated to their genomes) and the classification of EPSPS enzyme (resistant, sensitive or unclassified)**. (A) Series of boxplots show the absence of correlation between MAGs abundance in Phase II and their potential for antibiotic resistance. Each dot represents a MAG that is color-coded according to the potential resistance of their EPSPS. To facilitate visualization, a small amount of random variation (jitter) was added so dots would not overlap. Table 2 reports statistics of a linear model that tested how MAG abundance in Phase II controls could be explained by EPSPS classification, antibiotic resistance potential and MAG abundance in Phase I. (B) Regression tree with MAG abundance in controls of Phase II as the response variable and the following predictors: the EPSPS enzyme classification, the number of ARGs classified as antibiotic efflux, antibiotic inactivation or target alteration, and the MAG relative abundance in Phase I.

